# HBV enveloped particle secretion is positively regulated by GRP78 through direct interaction with preS1

**DOI:** 10.1101/2021.12.08.471876

**Authors:** Yueyuan Shi, Xin Jin, Shuang Wu, Junye Liu, Hongpeng Zhang, Xuefei Cai, Yuan Yang, Xiang Zhang, Jie Wei, Hui Peng, Miao Luo, Hua Zhou, Huihao Zhou, Ailong Huang, Deqiang Wang

**Affiliations:** The Key Laboratory of Molecular Biology of Infectious Diseases designated by the Chinese Ministry of Education, Chongqing Medical University, Yuzhong, Chongqing, 400016, China; College of Laboratory Medicine, Chongqing Medical University, Yuzhong, Chongqing, 400016, China; Division of Gastroenterology, Cedars-Sinai Medical Center, Los Angeles, California. Davis Bldg., Room 3094, 8700 Beverly Blvd., Los Angeles, CA 90048; Department of Clinical Laboratory, Yubei District people’s hospital, Yubei, Chongqing, 401120, China; Department of Clinical Laboratory, The Second Affiliated Hospital of Chongqing Medical University, Yuzhong, Chongqing, 400010, China; Clinical Laboratory, The Second Hospital of Harbin, Harbin City, Heilongjiang Province, 150056, China; Clinical Laboratory, The Affoliated Children Hospital of Xi’an Jiaotong University, Xi’an City, Shanxi Province, 710003, China; Clinical Laboratory, Honghui Hospital, Xi’an Jiaotong University, Xi’an City, Shanxi Province, 710054, China; Department of Clinical Laboratory, Chongqing Health Center for Women and Children, Yubei, Chongqing, 401132, China; Research Center for Drug Discovery, School of Pharmaceutical Sciences, Sun Yat-sen University, Guangzhou, 510006, China

**Keywords:** Hepatitis B virus, particle, secretion, GRP78, peptide, antiviral agent

## Abstract

Hepatitis B virus (HBV) infection is a common cause of liver diseases worldwide. Existing drugs do not effectively eliminate HBV from infected hepatocytes; thus, novel curative therapies are needed. Enveloped-particle secretion is a key but poorly studied aspect of the viral life cycle. Here, we report that GRP78 positively regulates HBV enveloped-particle secretion. GRP78 is the specific target of preS1 binding; HBV can upregulate GRP78 in liver cell lines and sera from chronic hepatitis B patients. GRP78 promoted intact HBV-particle secretion in liver cell lines and an HBV transgenic-mouse model. Some peptides screened from preS1 via phage display could inhibit viral-particle secretion by interacting with GRP78 via hydrogen bonds and hydrophobic interactions, thereby disturbing the interaction with HBV particles. These results provide insight into enveloped-particle secretion in the HBV life cycle. GRP78 might be a potential target for HBV-infection treatment via restricting GRP78–preS1 interactions to block viral-particle secretion.

**IMPORTANCE:** HBV is a major human pathogen. The virus More than 250 million n individuals globally are chronically infected with HBV and can result in the infected personsabout 800,000 die for HBV-related disease annually. The mature HBV enveloped particles containing HBs and the relaxed circular (RC) DNA genome could secreted extracellularly as virions. And these enveloped virions serves as the infectious HBV particles for initiating a complete life cycle of the virus. Here, we show the GRP78 is the specific target of preS1 binding, and it could positive regulate HBV enveloped particle secretion in cell and HBV transgenic mice model. Furthermore, several peptides screened from preS1 and Phage screening could inhibit viral particle secretion by interacting with GRP78 and disturbing its interaction with HBV, and then block the life cycle of the virus. This has important implications for our understanding of the mechanisms of antivirals that target intact HBV particles secretion.

---

Hepatitis B virus (HBV) is a major human pathogen, and more than 250 million people worldwide are chronically infected with HBV (1,2). Although a prophylactic vaccine is currently being used clinically, vaccinations are not effectively administered, and new infections continue to occur. The current care for chronic HBV infection, including six nucleos(t)ide and pegylated interferon-alpha, can significantly reduce viral load and prevent liver disease progression. Nucleoside analogs are currently the most potent drugs for inhibiting HBV; these analogs are also associated with drug resistance and require lifelong administration, as they do not effectively eradicate the virus from infected hepatocytes (2,3). Therefore, there is an urgent need to develop novel therapies for curing HBV infection (4,5).

As an enveloped virus, HBV contains three viral surface proteins: large hepatitis B virus surface protein, middle hepatitis B virus surface protein, and small hepatitis B virus surface protein. These proteins are integral membrane proteins that are embedded in the membrane surrounding the nucleocapsid. The nucleocapsid is assembled by the core protein (HBc) and harbors the viral DNA genome. A major characteristic of HBV is the secretion of various complete and incomplete viral particles, including nucleocapsids with or without viral nucleic acids, subviral particles (HBsAg) containing envelop proteins only, and virions containing an outer envelope enclosing an inner capsid with or without viral nucleic acids (6,7). In particular, a complete HBV particle contains an outer envelope enclosing an inner capsid with RC DNA genome content, which serves as the infectious HBV particle for initiating a complete viral life cycle. Consequently, the processes of assembly and transport in the cytoplasm and particle secretion are potential targets for antiviral drugs (8,9). Host factors can interact with HBc and envelope proteins to participate in HBV-particle secretion. Vps4 and the molecular endosomal sorting complexes required for transport (ESCRT) machinery could cooperate in the secretion of the HBV virion (10,11). Cellular kinases can regulate virion formation and promote intracellular capsid accumulation by modifying capsid phosphorylation (12–14). However, the exact nature of complete viral-particle secretion remains unclear.

Glucose-regulated protein 78 (GRP78) is an important endoplasmic reticulum (ER) chaperone that functions as a vital component in many cellular processes, including protein assembly, folding, and translocation across the ER membrane. The chaperone consists of two functional domains, including the amino terminal ATPase domain (nucleotide-binding domain [NBD], residues 31-407) and a carboxyl terminal peptide-binding domain (substrate-binding domain [SBD], residues 423-654) (15–18). GRP78 plays regulatory roles in the process and function of some viral envelope proteins, such as Middle East respiratory syndrome coronavirus (MERS-CoV), Ebola virus (EBOV), Zika virus, Dengue virus, Japanese encephalitis virus (JEV), Sindbis virus, hepatitis C virus (HCV), vesicular stomatitis virus, and influenza A virus. As such, GRP78 could serve as a receptor or cofactor to aid viral entry into host cells (19,20). Shurtleff et al. found that epigallocatechin gallate, which inhibits the ATPase activity of GRP78, can inhibit EBOV transcription and infection (21). Nain et al. also found that GRP78 plays an important role in JEV replication (22). Additionally, few studies noted that GRP78 might exhibit antiviral factors by inhibiting HBV infection; however, there is no direct evidence to clarify the antiviral effect of GRP78 (23–25).

In the present study, preS1 was used as the bait protein to screen for its binding protein by mass spectrometry analysis and GRP78 was identified as the corresponding binding protein. Further studies found that HBV could upregulate GRP78 in liver cell lines and chronic hepatitis B (CHB) patients. Transcriptome sequencing revealed that GRP78 could promote the transport and secretion of proteins in HBV-expression cell lines. Furthermore, GRP78 could promote the secretion of intact HBV particles in both the cell line and HBV transgenic mouse model. These results suggest that the host protein could positively regulate viral particle excretion by interacting with the preS1 of the HBV envelope protein. Importantly, some peptides, designed from preS1 or screening by phage display, could restrict viral particle secretion by interacting with the host protein to disturb the interaction between GRP78 and HBV particles. Our research provides new ideas for more in-depth study of HBV-particle secretion and for the use of GRP78 as a novel target for the development of antiviral therapy.

## RESULTS

### Screening host proteins interacting with preS1

The preS1 domain is localized on the outer membrane of HBV, and it interacts with host proteins, such as NTCP, which is the receptor of HBV (26). Therefore, the recombinant preS1-GST protein was prepared, and the fused GST tag with the bait protein facilitated subsequent affinity mass spectrometry experiments. The experimental group combined the biotin-labeled HepG2 hepatocyte lysate with the preS1-GST protein, and the control group was treated with biotin-labeled lysates of HEK293 cells and TM-GST irrelevant proteins to remove nonspecific signals. The results showed that some specific proteins with a molecular weight of 97-116 kDa were present in the HepG2 cell lysate (Fig. S1). Four proteins were identified in the preS1-GST protein captured from the HepG2 cell lysate by mass spectrometry, including keratin type II skeleton 1, type I skeleton 9, LanC-like protein 1 (LanCL1), and GRP78 (Fig. S1). The two skeleton proteins were present in relatively low quantities, and their presence was considered to be caused by contamination stochastically. LanCL1, a peptide-modifying enzyme component in eukaryotic cells, serves as a glutathione transferase and participates in breast cancer and prostate cancer (27). GRP78 could be galactosylated at three residues and might result in increased molecular weight, which further confirmed previous findings (28,29). Additionally, GRP78 is localized in the lumen of the ER, is involved in viral entry into the host cell, and/or regulates the replication of viruses, such as MERS-CoV, JEV, EBOV, HCV, and HBV (19,20,30). Confocal immunofluorescence detection revealed that both GRP78 and preS1 were spatially colocalized and mainly distributed in the cytoplasm (Fig. 1A). Co-immunoprecipitation revealed that GRP78 and preS1 could capture each other in HepG2 cells (Fig. 1B). To further verify the interaction between GRP78 and preS1, three GST-preS1 truncated proteins, including preS1-p1 (residues 1-65), preS1-p2 (residues 25-90) and preS1-p3 (residues 66-119), were expressed and purified, and these proteins were found to interact with GRP78 directly and with high affinity (K_d_ = 899±18.8 nM, 1,200±51.3 nM, and 431±20.1 nM, respectively) using a microscale thermophoresis assay (Fig. S2). These results imply that preS1 had multiple sites for interaction with GRP78. Consequently, it is worthwhile to further explore the interaction between GRP78 and preS1 in the HBV life cycle.

**Figure 1.**
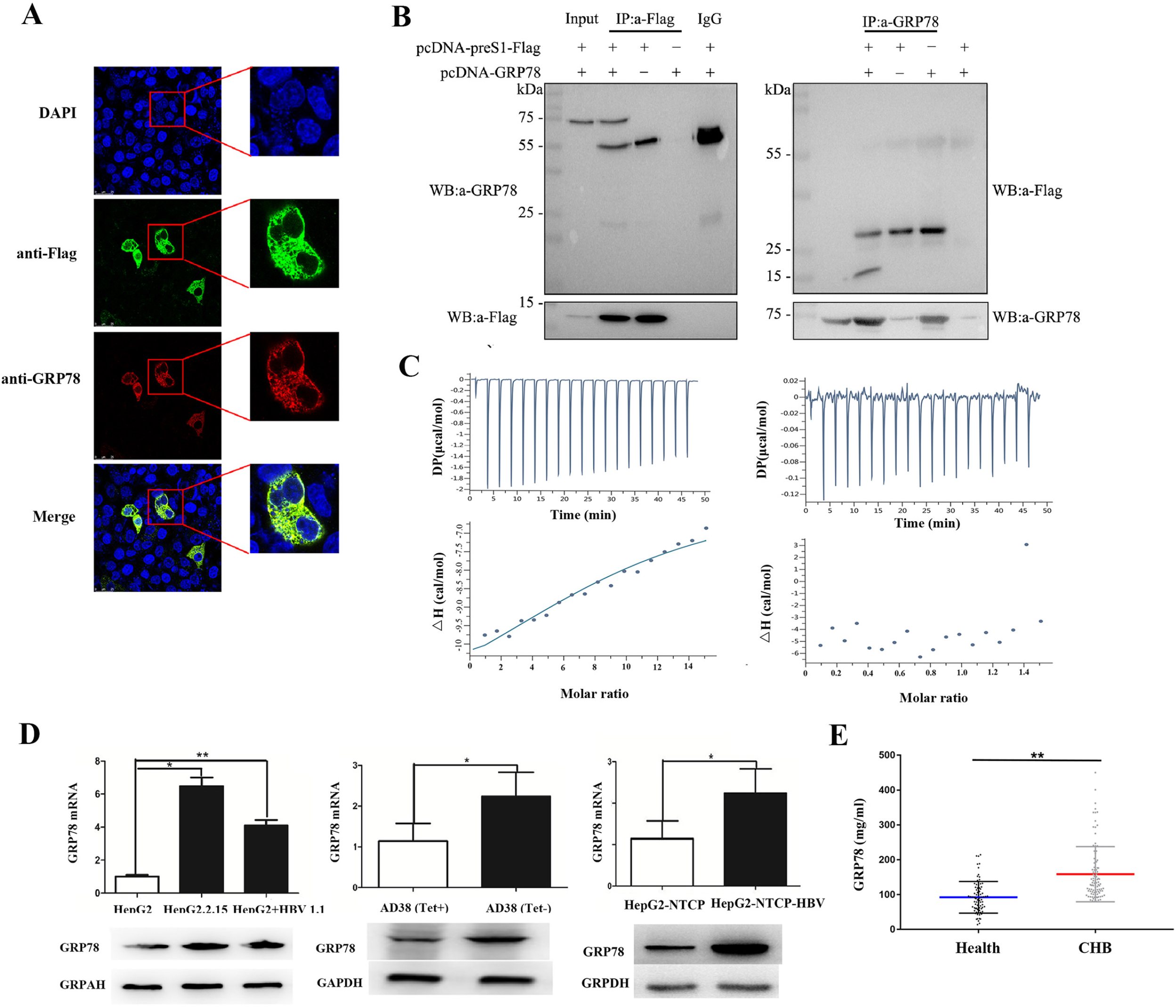
Direct interaction between preS1 of hepatitis B virus (HBV) and GRP78. **A.** Immunofluorescence confocal detection showed colocalization of GRP78 and preS1. Rabbit anti-GRP78 and mouse anti-preS1 were used to label two proteins in HepG2.2.15 cells that could stably express HBV, and these two proteins were mainly distributed in the cytoplasm. **B.** Immunoprecipitation was performed using agarose beads, and western blot detection confirmed that GRP78 was a specific binding target of preS1. **C.** The purified GRP78 protein could interact with HBV particles and adenovirus according to isothermal titration calorimetry (ITC). HBV and adenovirus particles were collected in the HepAD38 and 293 cell lines, respectively. **D.** HBV upregulated the expression of GRP78 in the HepG2, HepAD 38, and HepG2-NTCP cell lines. **E.** The GRP78-specific ELISA kit was used for quantitative detection, and unpaired *t*-test was used for statistical analysis.

Further analysis was conducted to determine the effect of HBV on GRP78. Both the mRNA and protein levels of GRP78 were significantly higher (over two folds) in HepG2.2.15 and HepG2 cells transfected with HBV plasmids than in HBV-negative cells. Similar changes were observed in HepAD38 cells treated with or without tetracycline (Fig. 1D). Additionally, infecting HepG2-NTCP cells with HBV increased the expression of GRP78 (Fig. 1D and S3). To further investigate the relationship between GRP78 and HBV, GRP78 serum levels were detected in 171 clinical serum samples from patients with CHB (Table S1). The experiments showed that the average GRP78 concentration (125.1 mg/mL) of chronic HBV carriers (n = 92) was higher than that in the serum of healthy individuals (92.31 mg/mL, n = 79) (Fig 1E). Detection of unfolded protein response molecular markers demonstrated that in addition to GRP78 and PERK, the expression of eIF2α, ATF4, IRE1α, and XBP1 were consistently elevated upon HBV overexpression in HepG2 cells (Fig. S3). These findings indicate that HBV expression could activate the PERK and IRE1 pathways to induce the ER stress response (31,32). Additionally, the purified recombinant GRP78 protein (Fig. S2) and HBV particles from the cell supernatant were collected and separated by ultracentrifugation at 4 °C for ITC experiments. The experiments revealed that GRP78 could interact with HBV particles with a dissociation constant K_d_ = 75.6×10^-6^ M (Fig. 1C), whereas there was little interaction between GRP78 and adenovirus. Together, the above experiments verified that HBV could promote GRP78 expression, and GRP78 could directly interact with both preS1 and HBV particles.

### GRP78 promotes HBV secretion in cell supernatant

preS1 is the viral binding site for NTCP, the receptor of HBV, and is the unique domain at the outermost surface of the HBV particle. However, GRP78 binding to preS1 had little effect on viral entry into host liver cells (data not shown). To screen the function of the interaction between GRP78 and preS1, an adenovirus overexpressing GRP78 (pAd-GRP78) was constructed and used to infect HBV-expressing cells. The effect of pAd-GRP78 overexpression was verified by RT-qPCR and western blot analysis of GRP78 mRNA and protein levels in both HepG2.2.15 and HepAD38 cells (Fig. S4). Next, the transcriptome sequencing experiments in HepG2.2.15 infected with pAd-GRP78 showed that multiple pathways of gene transcription (including DNA replication, RNA degradation, and RNA transport) and cancer-related roles, such as p53 signaling and cell cycle, were activated. The most activated pathway included the proteins involved in protein export and transshipment (Fig. S5). Consequently, these data provide information that may allow us to determine which secretion pathway was activated by GRP78.

According to the components of HBV particles, anti-HBc and anti-preS1 could be used to detect naked core (NC) particles and intact virus particles, respectively. HepG2.2.15 cells were infected with pAd-GRP78 and pAd-GFP to determine the effect of the host chaperone protein. The particle gel assay showed that GRP78 had little effect on both the core and enveloped particles in the cytoplasm and similar core granules in the cell supernatant, whereas it significantly increased particles with an envelope in the supernatant (Fig. 2A). Additionally, according to Southern and western blot, levels of both the viral DNA and envelope protein were much higher in the supernatant of pAd-GRP78-treated HepG2.2.15 cells than in that of cells infected with pAd-GFP (Fig. 2B). Furthermore, more intact viral particles were detected in the supernatant of HepAD38 cells treated with Ad-GRP78 than in that of cells treated with Ad-GFP (Fig. 2D). However, according to ELISA, both HBsAg and HBeAg showed little change in the two cell lines with GRP78 overexpression (Fig. 2C and 2E). These results suggest that GRP78 overexpression could promote the secretion of intact virions and exhibited little effect on both free antigen and core particles.

**Figure 2.**
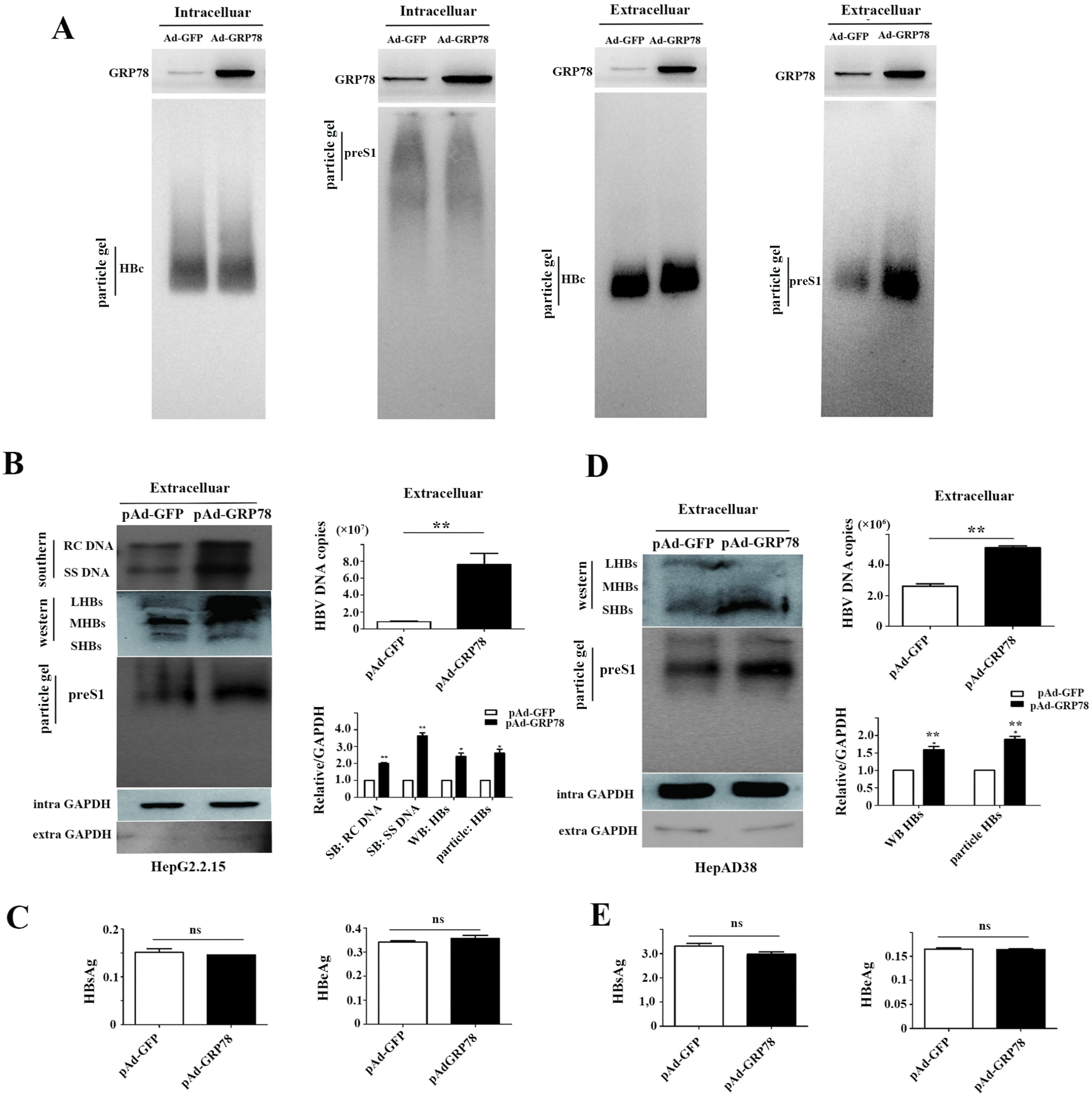
Overexpression of GRP78 promoted HBV-particle secretion. **A.** Particle gel assay to detect the HBV particles (core particle and intact viral particle) in the supernatant and cytoplasm of HepG2.2.15 cells infected by pAd-GRP78. **B.** Overexpressing GRP78 promoted HBV-particle secretion in HepG2.2.15 cells infected with GRP78Ad. HBV DNA was subjected to real-time quantitative polymerase chain reaction (RT-qPCR) and Southern blotting. The HBV particles were detected by viral particle gel experiments with preS1 antibody. **C.** The relative level of HBV e antigen (HBeAg) and HBV surface antigen (HBsAg) in cellular supernatant of HepG2.2.15 treated with GRP78Ad were subjected to ELISA. **D.** Overexpressing GRP78 promoted HBV secretion extracted from HepAD38 cells infected with GRP78Ad. HBV DNA were subjected to RT-qPCR and Southern blotting. The HBV particles were detected by viral particle gel experiments with preS1 antibody. **E.** The relative levels of HBeAg and HBsAg in the supernatant of HepAD38 cells treated with GRP78Ad were observed via ELISA.

### GRP78 knockdown inhibits HBV secretion in cell supernatant

Next, interfering RNA was used to inhibit GRP78 expression in HepG2.2.15 and HepAD38 cells (Fig. S4). The results of Southern and western blot showed that both the number of extracellular HBV DNA copies and envelope proteins were decreased significantly. Furthermore, HBV enveloped particles were present at much lower levels in the culture fluid of siRNA-treated cells than in cells with normal GRP78 expression (Fig. 3A and 3B). However, according to ELISA, the secretion of HBsAg and HBeAg and knockdown of GRP78 expression did not reduce the concentration of the two antigens in the supernatant of the two cell lines (Fig. 3C and 3D). These results further suggest that GRP78 positively promotes HBV-particle secretion and has little effect on the free HBsAg in the serum.

**Figure 3.**
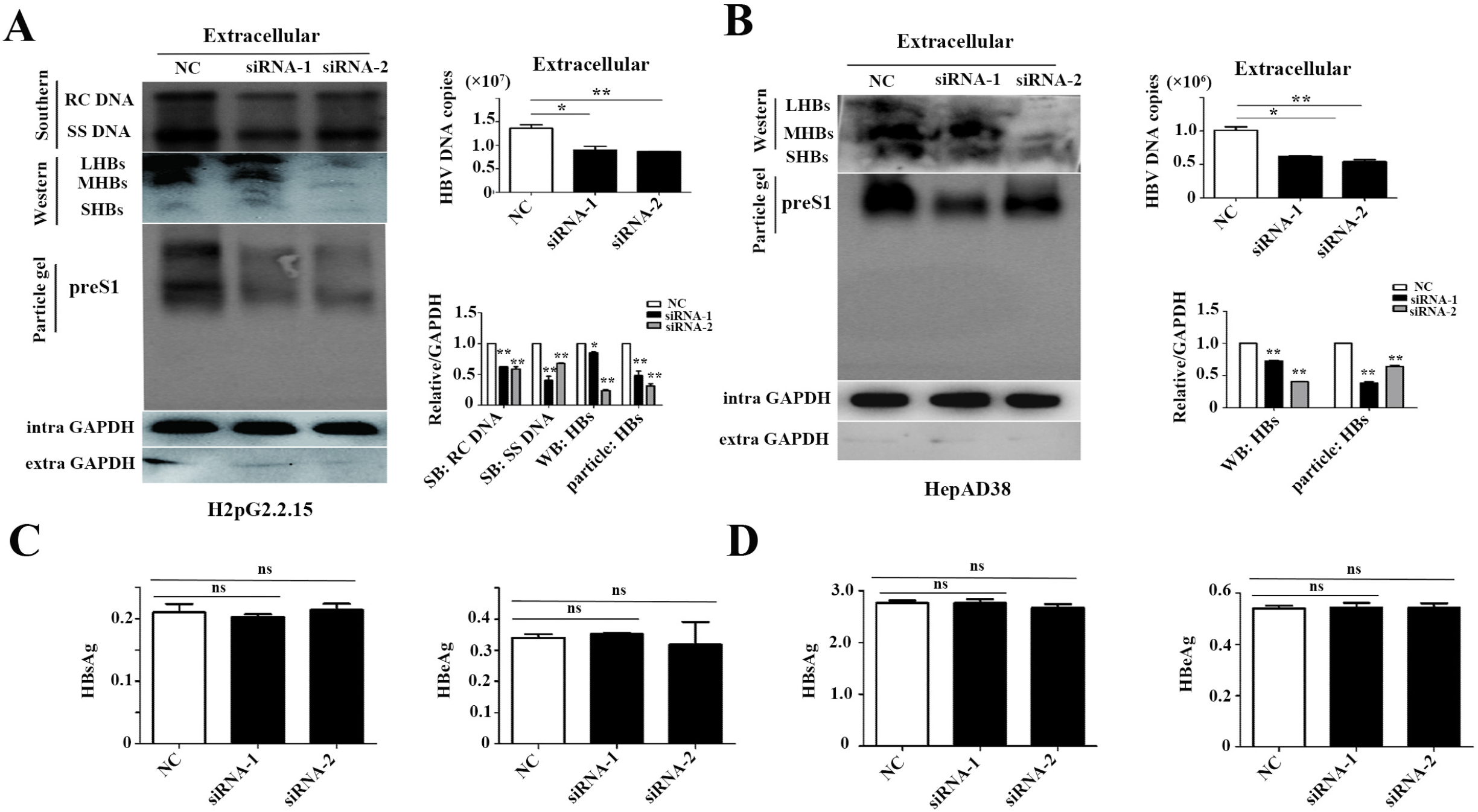
GRP78 knockdown in HepG2.2.15 and HepAD38 cells disturbed HBV-particle secretion. **A.** GRP78 knockdown inhibited HBV-particle secretion in HepG2.2.15 cells infected with GRP78 siRNA. HBV DNA was subjected to real-time quantitative polymerase chain reaction (RT-qPCR) and Southern blotting. The HBV particles were detected by viral particle gel experiments with preS1 antibody. **B.** The relative levels of HBeAg and HBsAg in the cellular supernatant of HepG2.2.15 cells treated with GRP78 siRNA were observed using ELISA. **C.** GRP78 knockdown promoted HBV secretion in HepAD38 cells infected with GRP78 siRNA. HBV DNA was subjected to RT-qPCR and Southern blotting. The HBV particles were detected by viral particle gel experiments with preS1 antibody. **D.** The relative levels of HBeAg and HBsAg in cellular supernatant of HepAD38 cells treated with GRP78 siRNA were observed using ELISA.

### GRP78 promotes HBV-particle secretion *in vivo*

To further investigate whether HBV particles are affected by GRP78 *in vivo*, eight HBV transgenic mice (HBV TgM) were randomly divided into control and experimental groups, which were treated with pAd-GRP78 or pAd-GFP by tail vein injection (Fig. 4A). Three weeks later, we extracted serum and liver tissue from all experimental transgenic mice according to animal ethics guidelines. Both western blot and immunohistochemical experiments showed that GRP78 was overexpressed in the liver tissues of HBV TgM treated with pAd-GRP78 (Fig. 4B and 4C). The results of particle gel assay and RT-qPCR showed that GRP78 significantly upregulated the HBV DNA copy number and intact viral particles in the serum of HBV TgM infected with pAd-GRP78 (Fig. 4B and 4D). However, there was little difference in the serum HBsAg and HBeAg levels between the two groups. These findings are consistent with the results of previous *in vitro* experiments, which show that GRP78 overexpression could promote HBV-particle secretion. The *in vivo* experiments further confirmed that GRP78 could promote the secretion of intact viral particles and had little effect on both HBeAg and HBsAg.

**Figure 4.**
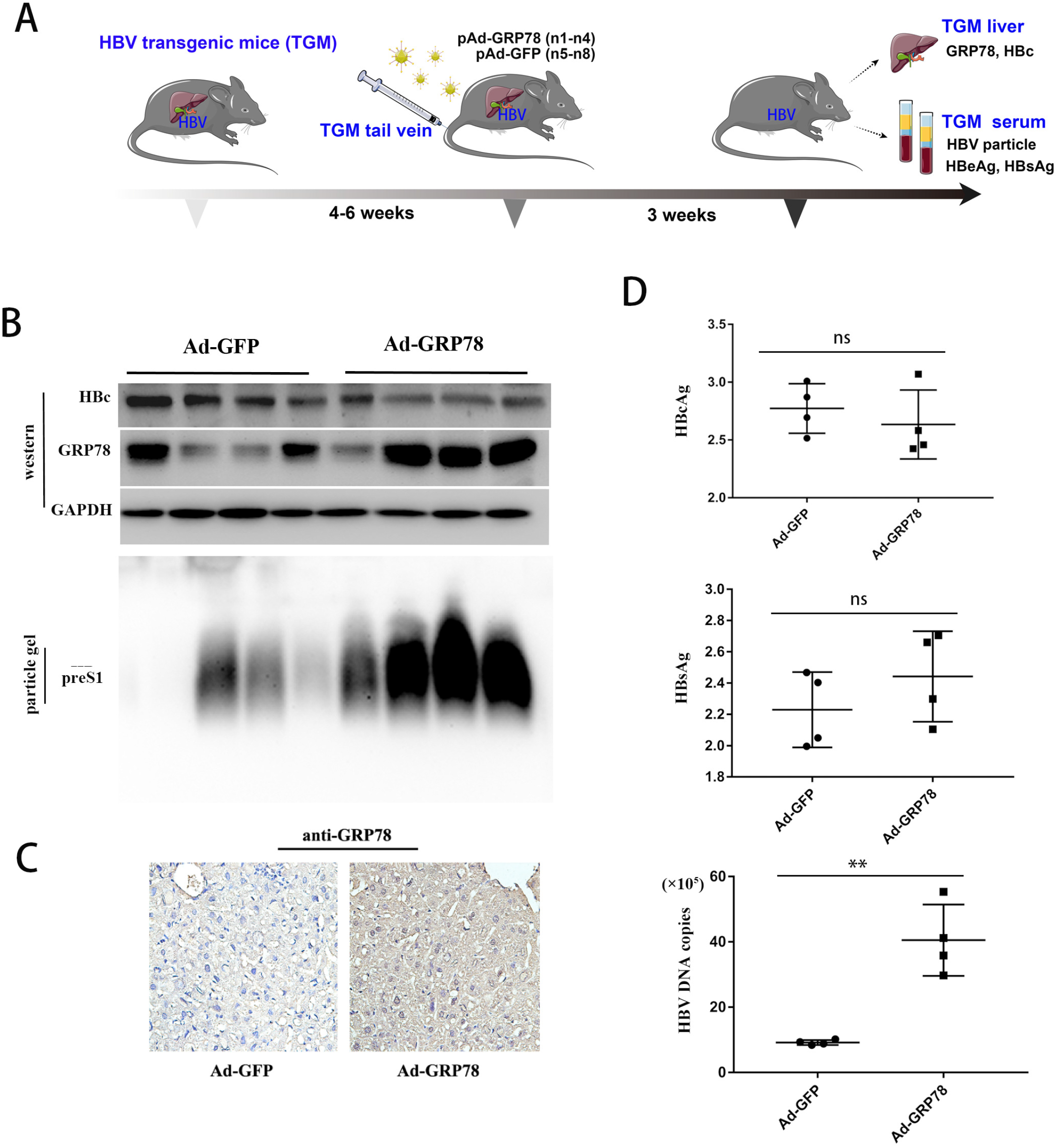
GRP78 overexpression in HBV transgenic mice (TGM) increased viral particle secretion. **A.** The mice (4-6 weeks) were randomly allocated to two groups (n = 4 per group). We dissolved 7×10^9^ GFU GRP78Ad (n1-4) or GFPAd (n5-8) in 0.3 mL 0.9% normal saline and injected it into mice through the tail vein. The mice were sacrificed at three weeks after injection, and the liver tissue and serum were collected. **B.** GRP78 expression in liver tissue lysates was tested by western blotting with GAPDH as an internal control. The HBV particles from serum were detected by viral particle gel experiments with preS1 antibody. The serum of the mice was collected to detect the copy number of HBV DNA and the secretion level of HBV virions. **C.** Immunohistochemistry analysis detected less HBc in sectioned liver samples in the presence of TGM treated with GRP78Ad or GFPAd at 1 dpi. **D–E.** The amount of HBeAg, HBsAg and HBV DNA before and after adenovirus injection and TGM serum extraction was analyzed by ELISA or qPCR, respectively.

### SBD and NBD of GRP78 participate in HBV-particle secretion

GRP78 is an important molecular chaperone protein of *Homo sapiens;* it can be secreted into the extracellular space and serum because of its signal peptide sequences (Fig. 5A) (33,34). Adenovirus (pAd-GRP78-Δss) expressing the truncated GRP78 signal peptide was constructed (Fig. S6). Both particle gel experiments and viral copies in the supernatant showed that overexpression of both GRP78-Δss and GRP78 could increase the concentration of viral particles to the same degree in the HepH2.2.15 cell line (Fig. 5B). The background expression of the GRP78 gene could not be removed because knockout of this gene would lead to cell death (data not shown). Although removal of the signal peptide should result in the prevention of GRP78 secretion, it does seem that the signal peptide does not affect the promotion of virus-particle secretion by GRP78. The SBD and NBD of GRP78 were arranged in a remote region with a flexible linker sequence, suggesting that the effect on viral particle secretion might be undertaken by a single domain (33,34). The SBD and NBD were overexpressed in the HepH2.2.15 cell line upon infection with pAd-SBD and pAd-NBD (Fig. S6). Meanwhile, both the particle gel and qPCR assays revealed that both these domains exhibited little effect on viral secretion and HBV copies in the cell-culture fluid, which may be due to the high background expression of GRP78 homoplastically (Fig. 5C). Additionally, according to the three-dimensional structure, adenovirus overexpressing five active site mutants (including NBD mutants [T38A and T229A] and SBD mutants [F451A, I463A, and R492A]) was constructed and used to infect HepG2.2.15 cells (Fig. S6) (33,34). The experiments showed that these five mutants exhibited a much lower ability to promote HBV-particle secretion compared to wild-type GRP78. In particular, T38A, F451A, I463A, and R492A almost lost their ability to promote viral particle secretion, while the T229A mutant retained 50% of the effect (Fig. 6D). The above data suggest that the SBD and NBD might be involved in collaboratively promoting viral particle secretion, implying that inhibitors of GRP78 might obstruct HBV secretion.

**Figure 5.**
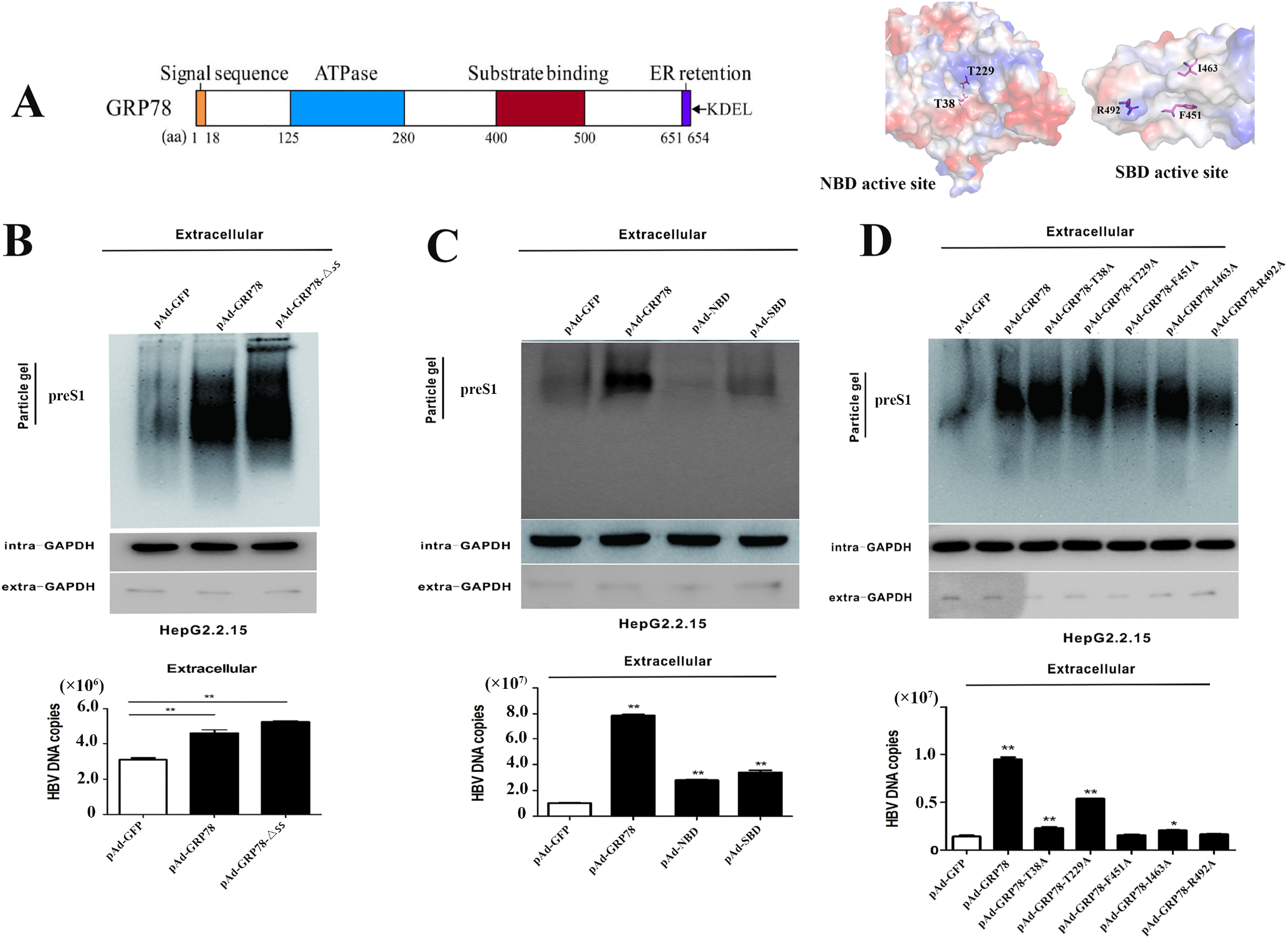
Domains of GRP78 participate in HBV-particle secretion. **A.** Schematic showing the GRP78 sequence and domains. The signal sequence, ATPase domain, and substrate binding domain are highlighted in yellow, blue, and aubergine boxes, respectively. **B.** The HBV particles in HepG2.2.15 cells treated with pAd-GFP, pAd-GRP78, and pAd-GFP-△ss were detected with preS1 antibody. **C.** The HBV particles in HepG2.2.15 cells treated with pAd-GFP, pAd-GRP78, pAd-NBD, and pAd-SBD were detected with preS1 antibody. **D.** The HBV particles in HepG2.2.15 cells treated with Ad-GFP, Ad-GRP78-T38A, Ad-GRP78-T229A, Ad-GRP78-F451A, Ad-GRP78-I463A, and Ad-GRP78-R492A were detected with preS1 antibody.

**Figure 6.**
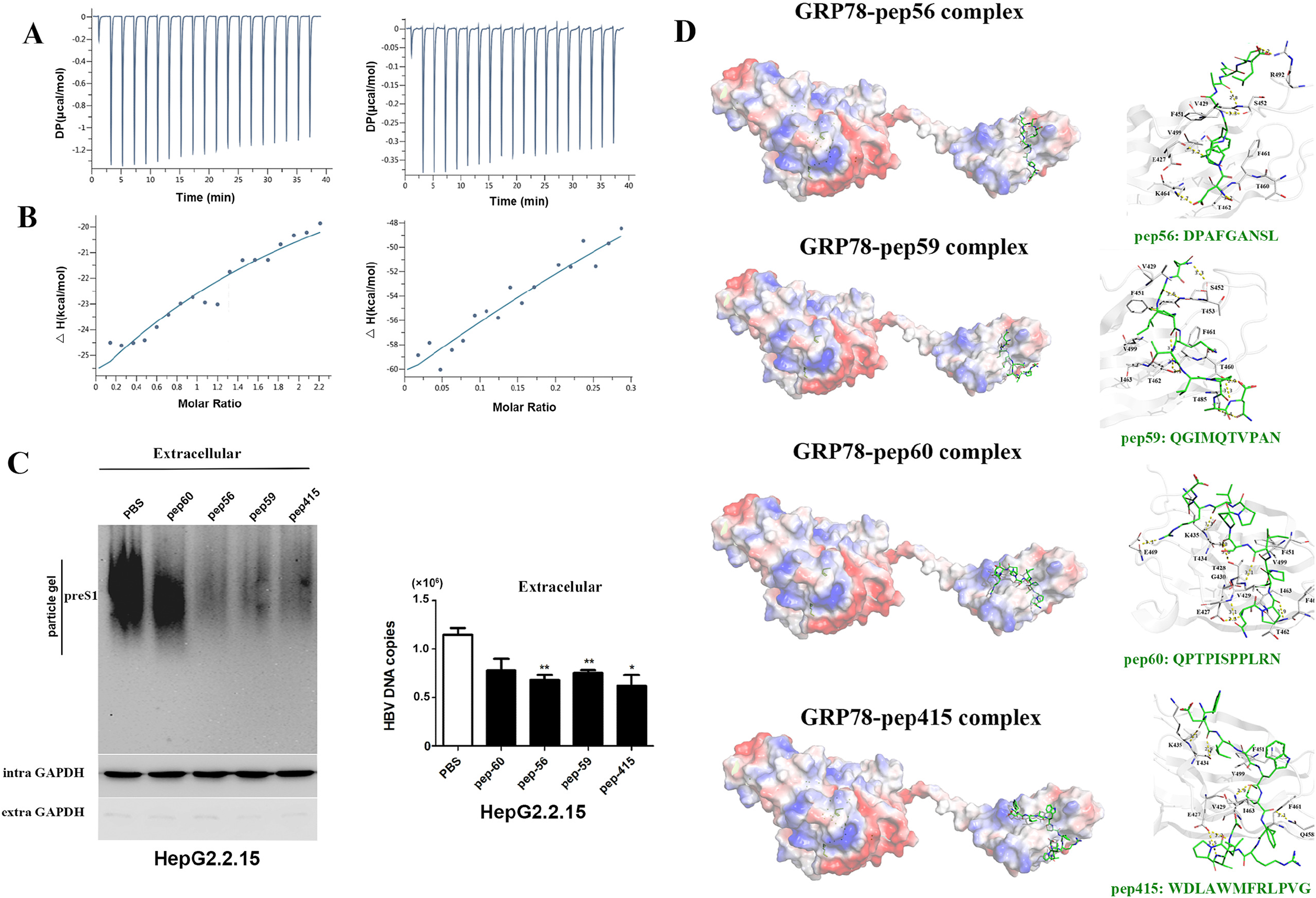
The preS1 peptide inhibited HBV-particle secretion. **A,B.** Interaction between recombinant GRP78 and peptide (pep56 and pep59) according to ITC assay. **C.** The HBV particles of HepG2.2.15 cells treated with pep56, pep59, pep60, and pep415 were detected with preS1-antibody. **D.** Complex structure model of GRP78 and the peptide complex. The GRP78 structure model was obtained from PDB (code: 6HAB. http://www.rcsb.org/structure/6HAB), and the complex models were calculated from AutoDock Vina (http://vina.scripps.edu/) (Table S2). H-bond interaction network between GRP78 and peptide (pepp56, pep59, pep60, and pep415).

### preS1 disturbs the promotion of HBV-particle secretion by GRP78

According to the structural and functional features of GRP78, the SBD binds to hydrophobic peptides, and the NBD provides the energy for the interaction by hydrolyzing ATP to ADP. Several molecules, including catechins (epigallocatechin gallate, honokiol, and aspirin, can restrain its ATPase activity, and some peptides can interact with the SBD directly (34). Considering the direct interaction of GRP78 with preS1 and its promotion of HBV-particle secretion, the peptide WDLAWMFRLPVG (pep145) was synthesized and directly interacted with recombinant GRP78 (Fig. 6B) (34,35). Compared with the control, pep145 inhibited 80% of the activity of GRP78, thereby positively regulating viral particle secretion (Fig. 7D). To further explore whether the effect of GRP78 on HBV is based on its binding to HBV preS1, hydrophobicity analysis of the amino acid sequence of preS1 was carried out using bioinformatics. Furthermore, three hydrophobic peptides, including DPAFGANSL (pep56), QGIMQTVPAN (pep59), and QPTPISPPLRN (pep60), were designed to block the interaction between preS1 and GRP78 (16). The MTS cell proliferation assay showed that the cytotoxicity of these three peptides was very low in HepG2.2.15 cells (data not shown). ITC assay showed that pep56 and pep59 interacted with recombinant GRP78 with a high affinity (K_d_ = 6.588×10^-9^ M and 1.057×10^-8^ M, respectively) (Fig. 7A), whereas there was little interaction between pep60 and GRP78 (data not shown). Interestingly, similar to pep145, both pep56 and pep59 inhibited the effect of GRP78 on HBV-particle secretion in HepG2.2.15 cells overexpressing GRP78; nevertheless, pep60 exhibited little effect on viral secretion (Fig. 6B). To further investigate the binding mode of GRP78, molecular docking of these peptide-GRP78 complexes was performed (Table S2). Molecular docking analysis revealed that pep415, pep56, pep59, and pep60 could bind with GRP78 and had good docking scores of −12.2496 kcal/Mol, −10.0518 kcal/Mol, −11.1134 kcal/Mol, and −11.3549 kcal/Mol, respectively. Though the hydrophilic interactions and array of H-bond interactions, it was coped by the strong hydrophobic interaction for the peptide between the GRP78 residues (F451, V499, I463, V429, and F461) and the hydrophobic residues located in the middle of the peptide, including Phe4 (pep56), Ile3 (pep59), and Met6 (pep415). However, there was much lower hydrophobic interaction between GRP78 and pep60; this finding may explain the weak interaction observed by ITC, leading to a poor effect on virus secretion inhibition (Fig. 6C).

**Figure 7.**
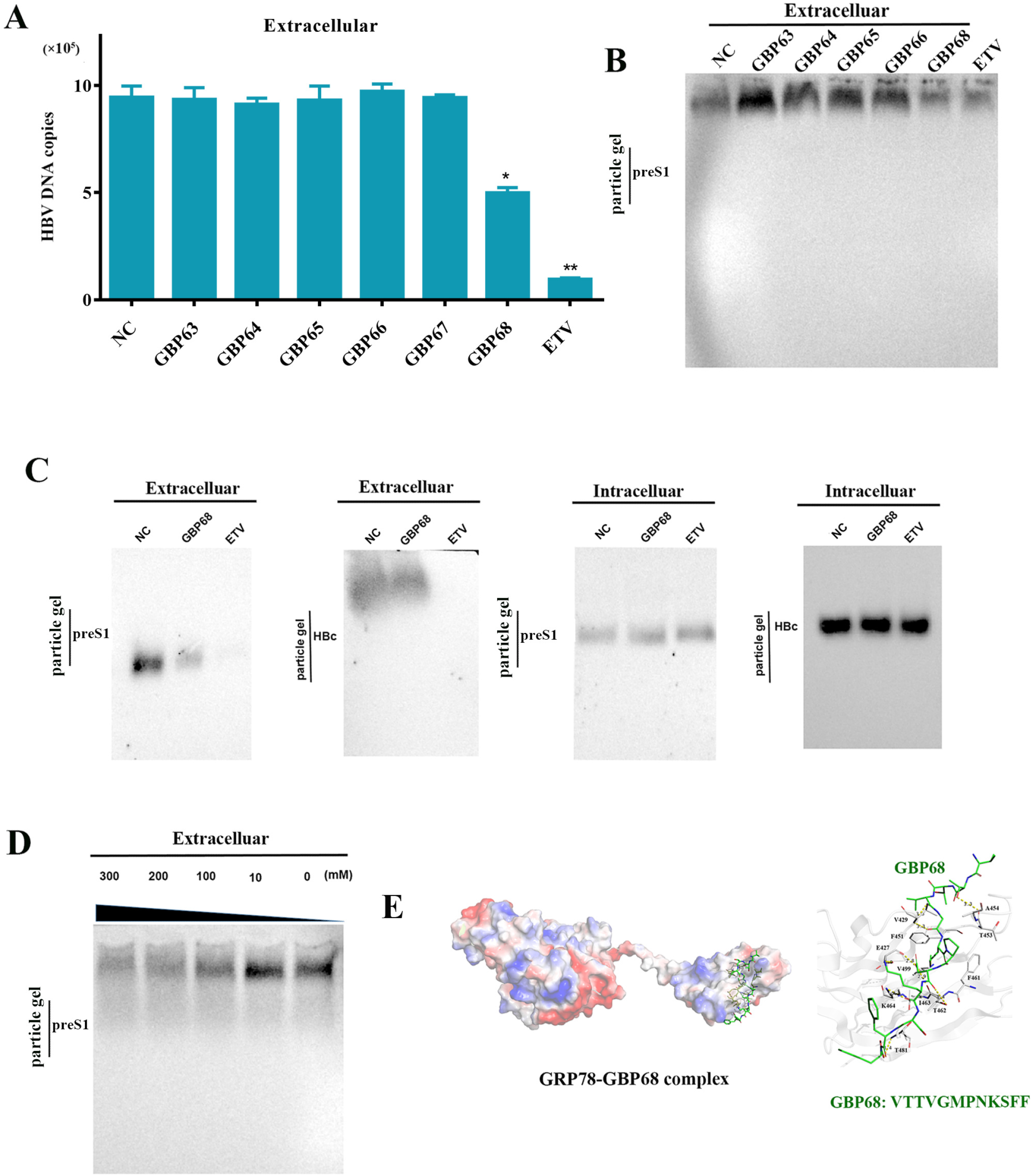
Peptides screened by phage display inhibit HBV-particle secretion. **A.** The HBV DNA copies in cellular supernatant from HepG2.2.15 cells treated with GBP63, GBP64, GBP65, GBP66, GBP67, GBP68, and entecavir (ETV). **B.** The HBV enveloped particles in cellular supernatant from HepG2.2.15 cells treated with GBP63, GBP64, GBP65, GBP66, GBP67, GBP68, and ETV, according to particle gel assay. **C.** The HBV particles (core particle and intact virus particle) in the supernatant and cytoplasm of HepG2.2.15 cells treated with GBP68 and ETV, respectively. **D.** The HBV enveloped particles secreted from HepG2.2.15 cells treated with different concentrations of GBP68. **E.** The complex structure model of the GRP78 and GBP68 complex. The GRP78 structure model was cited from PDB (code: 6HAB), and the complex models were calculated from AutoDock Vina (http://vina.scripps.edu/) (Table S2). **F.** The model of GRP78 regulates HBV-particle secretion. GRP78 promotes intact HBV-particle secretion by directly interacting with preS1 located at the viral envelope. Some peptides could inhibit HBV particle excretion by binding to GRP78 with high affinity, which in turn reduced the inhibitory interaction between GRP78 and preS1/HBV particles.

### Screening peptides to inhibit HBV-particle secretion

Phage display is a powerful method for screening binding peptides from a highly diverse combination of libraries (36). To design inhibitors based on the GRP78 protein and the preS1 binding protein, a phage 12-peptide library was utilized for screening candidate peptides. After three rounds of biopanning, 144 clones were selected for Sanger sequencing (data not shown). There were eight kinds of peptides with multiple frequencies among the 60 sequenced clones, including GBP 63 (pep7), GBP 64 (pep11), GBP 65 (pep16), GBP 66 (pep19), GBP 67 (pep32), and GBP 68 (pep51). Two peptides, pep9 and pep46, were found to be too hydrophobic to be synthesized (Table S3). The MTS cell proliferation assay showed that the six synthesized peptides exhibited little cytotoxicity to HepH2.2.15 cells (data not shown), suggesting that they did not affect host cell metabolism or viability and are safe for use in mammalian cells at concentrations as high as 200 μM. Additionally, among these synthesized molecules, five peptides could interact with the recombinant GRP78 with strong affinity (Table S4); meanwhile, ITC experiments showed that there was little interaction between GRP78 and GBP67. Consequently, the inhibitory activity of the other five peptides on HBV-particle secretion was determined by particle gel and HBV DNA assays. GBP68 exhibited a much stronger inhibitory effect than GBP63, GBP64, GBP65, and GBP66 in the supernatant of HepG2.2.15 cells (Fig. 7A and 7B). Furthermore, GBP68 inhibited HBV-particle secretion both in HepG2.2.15 and HepAD38 cells but exhibited little effect on viral particles in the cytoplasm and core particle excretion (Fig. 7C and S8). The inhibitory effect of HBV-particle secretion by GBP68 was dose-dependent (Fig. 7D). Figure 7E represents the best-scoring binding poses of GBP68 in complex with GRP78 (docking score of −12.008 kcal/M), as well as the key interacting residues at the binding pose. It was observed that the GBP68 generated nine hydrogen bonds within the range of 3.50 Å with GRP78, and the key residue, Met6, inserted into a hydrophobic bag formed by Phe451, Phe461, Val499 and Val511. There was also a strong hydrophobic effect between GBP68 and GRP78. Taken together, the above experiments further demonstrated that the interaction of peptides with GRP78 could inhibit HBV secretion (Fig. S9).

## DISCUSSION

The HBV life cycle includes a phase that occurs within the hepatocyte nucleus, wherein HBV DNA is converted to cccDNA, which is a highly stable double-stranded circular DNA structure. cccDNA serves as a template for viral RNA transcription and can persist indefinitely within the long-lived hepatocyte nucleus, providing a reservoir for viral replication. The secretion of complete HBV virions requires interactions between host proteins and viral envelope proteins; thus, inhibition of virus secretion could represent an efficient tool to prevent the spread of viral infection. However, exact regulatory mechanism for enveloped virion secretion is currently unclear. In this study, GRP78 interacted specifically with HBV particles via the viral envelope protein preS1, and HBV can upregulate GRP78 expression and activate the ERS system; conversely, GRP78 mainly acts on HBV virion secretion and promotes HBV-particle secretion in a nonreplicating manner. Moreover, the hydrophobic peptide preS1 interacts with GRP78 with strong affinity and inhibits HBV enveloped-particle secretion. Additionally, peptide screening by phage display showed that GBP68 could also interact with recombinant GRP78 and significantly inhibit viral secretion. To the best of our knowledge, this is the first study to provide new avenues for more in-depth study of HBV enveloped-particle secretion and new targets for antiviral drugs.

Viral particle secretion is a key factor in the HBV life cycle. Virion morphogenesis involves interactions between HBc, envelope proteins (HBsAg), and host factors, such as components of the ESCRT machinery. Vps4, a host factor known to be involved in cellular vacuolar protein sorting, is associated with ESCRT-mediated membrane dynamics. It has been reported that HBV replication and virion secretion are significantly inhibited by Vps4 (37,38). ESCRT can mediate the export of a number of viruses and over 15 ESCRT factors, including CHMP3, CHMP4B, CHMP4C, EAP20, EAP30 and EAP45, that are required for HBV replication and secretion (10,11,38,39). Host cellular kinases, including protein kinase C (PKC), SRPK1/2, CDK2, PLK1, and protein phosphatase 1, can regulate the phosphorylation of HBV capsids, resulting in the inhibition of virion formation and promotion of intracellular capsid accumulation (12–14). BST-2/tetherin, an IFN-inducible antiviral cellular protein, could restrict HBV virion secretion in HepG2 cells, but this effect was not observed in HuH-7 cells (40). These host factors exhibit some effect on HBV capsid and/or particle secretion, whereas there is little evidence that they could not directly interact with preS1; consequently, it has been suggested that these proteins participate in viral secretion indirectly. In this study, GRP78 directly interacted with both preS1 and HBV particles and positively regulated HBV-particle secretion. In addition, several host proteins other than GRP78 are involved in viral particle secretion; further research is needed to clarify whether these proteins collaborate with GRP78 to promote viral secretion.

Taken together with the current literature, these results allow us to propose a model for HBV enveloped-particle secretion that is positively regulated by the human host shock protein, GRP78 (Fig. S9). While additional biochemical and structural studies will be required to directly confirm the interactions predicted herein, evidence provided by our experiments using cell models and transgenic mice strongly support this model. When the mature nucleocapsids are further packed as intact viral particles by envelope proteins, the preS1 domain of HBV L protein is located at the outermost layer for host protein interaction. This domain then directs viral secretion. Consequently, the host protein that directly interacts with preS1 plays a key role in HBV-particle secretion. As such, we screened the preS1 binding protein in human liver cell lines. The γ2-adaptin HEAD domain (amino terminal) can bind HBV nucleocapsids and mediate virion export by the host ESCRT machinery, and its EAR domain (carboxyl terminal) interacts with preS1 and participates in HBV replication (41). NTCP specifically interacts with the preS1 region of the HBV L protein and serves as an entry receptor for HBV infection (27,42). Additionally, GRP78 could bind to the preS1 region of HBV and function as an intracellular antiviral factor by inhibiting HBV infection; however, there is no clear evidence of the mechanism of the antiviral effect of this host protein. Using affinity capture and high-end mass spectrometry analysis with the established *Treeshrew* transcriptome database, we further revealed that GRP78 specifically interacts with the preS1 region. Additionally, the host protein exhibited little effect on HBV entry into human liver cells. GRP78 could promote the secretion of HBV enveloped particles in the cellular supernatant and serum of transgenic mice by improving the virus concentration in the cytoplasm. These results suggest that GRP78 could positively regulate HBV-particle secretion both *in vitro* and *in vivo*. Consequently, GRP78 may provide a target for antiviral drugs to inhibit HBV-particle secretion.

According to our experimental results, GRP78 could bind to both preS1 and HBV by its SBD at the carboxyl terminus. Thus, we hypothesized that if a peptide interacts with GRP78, it should be able to block or interfere with its binding to the viral envelope protein, thereby blocking or inhibiting viral secretion. Indeed, we identified fragments of preS1 that could interact with GRP78 directly, thereby inhibiting enveloped-particle secretion. Similarly, we used phage display technology to screen for binding peptides of GRP78, which can also inhibit viral secretion. These results further confirmed that GRP78 positively regulates viral secretion by binding to the envelope protein preS1. If we can screen for peptides interacting with GRP78 with sufficient affinity, it may be possible to develop antiviral drugs that block viral secretion.

As a molecular chaperone, GRP78 plays an essential role in the development of envelope proteins of some viruses, such as Sindbis virus, Hepatitis C virus, Vesicular Somatitis virus, and Influenza A virus. GRP78 functions as an attachment factor for binding to the MERS-CoV spike protein and enhances viral entry into host cells (19). Data derived mainly from the study of other coronaviruses provide sufficient evidence to discuss the putative role of CD147 and GRP78 as entry receptors for SARS-CoV-2. Additionally, the spike protein of SARS-CoV-2 could stably interact with the head of GRP78, implying that it might be involved in the entry of SARS-CoV-2 (43). Coxsackievirus A9 (CAV9) cloud binds to the host receptor GRP78 and integrin αvβ3, and then uses MHC-I for entry into host cells (44). GRP78 functions as a receptor for JEV by directly interacting with envelope domain III (ED3) (45). GRP78 interacts with the envelope and has been identified as a receptor for dengue virus serotype 2 (46). Similarly, GRP78 could interact with the HBV envelope protein; however, this host protein does not exhibit an effect as a receptor to aid HBV entry into host cells. Instead, GRP78 is involved in viral particle secretion, which provides some new perspectives for clarifying the function of GRP78 in the viral life cycle.

In the present study, GRP78 was found to interact with preS1 directly, and it positively regulated HBV-particle secretion. The peptide inhibited enveloped-particle secretion by disturbing the interaction between GRP78 and preS1. These findings provide a basis for more in-depth study of HBV enveloped-particle secretion, and GRP78 may function as novel target for designing antiviral therapy through the inhibition of HBV secretion.

## MATERIALS AND METHODS

### Cell culture, plasmids, and mice

Human hepatoma cell lines HepG2, HepG2.2.15 and HepAD38 were saved by our research group. HepG2.2.15 cells were cultured in Dulbecco’s modified Eagle medium containing 500 μg/mL G418, and the remaining cells were cultured in Dulbecco’s modified Eagle medium. All cells were supplemented with 10% (v/v) fetal calf serum (FCS) (ABI) at 37 °C in 5% CO_2_. HBV replication plasmids, pcDNA3.1-HBV1.1, and the recombinant plasmids expressing GRP78 and preS1 were all preserved in our laboratory. Adenoviruses expressing GFP, wild types GRP78 and its mutants were constructed and prepared by our group. HBV transgenic C57BL/6J male mice were gifted by Prof. NX Shao (Xiamen University), and the breeding and experimental studies involving transgenic mice were carried out at the Experimental Animal Center of Chongqing Medical University.

### Patients

Table S1 shows the characteristics of the patients in the present study, including 103 women and 68 men with a median age of 55.2 (18–65 years). The study protocol was approved by the Research Ethics Committee of Chongqing Medical University and written informed consent was obtained from each patient.

### Particle gel assay

The HBV virion was identified by the particle gel assay (Ran et al., 2017). HBV-producing cells were cultured, and the cell culture medium was harvested. The cell supernatant was added to 35% PEG8000 and rotated at 4 °C for more than 2 h. The viral particles were dissolved in TNE buffer and incubated overnight at 4 °C. The dissolved viral particles were subjected to agarose gel electrophoresis. Then, viral particles were transferred to a nitrocellulose membrane, maldehyded for 10 min, methanoled for 30 min, blocked with 5% nonfat milk, and incubated with anti-preS1 (for intact particles) or anti-HBc (for capsid particles) for 1 h at RT. After incubation with anti-mouse secondary antibody (1:10,000) at 4 °C overnight, the membrane was visualized using the ECL Western Blotting Detection Kit (Bio-Rad, USA).

### RNA extraction and real-time quantitative PCR

Total RNA was extracted from cells (or liver tissues from HBV transgenic mice) using TRIzol reagent (Invitrogen, USA). First-strand cDNA was synthesized as previously described. RT-qPCR was performed using a Bio-Rad sequence detection system according to the manufacturer’s instructions using a double-stranded DNA-specific SYBR Green Premix Ex TaqTM II Kit (Roche, Switzerland). Experiments were conducted in duplicate in three independent assays. Relative transcriptional fold was calculated as 2-ΔΔCT. GAPDH was used as the internal control for normalization.

### Protein extraction and western blot detection

Total protein lysates were extracted from hepatoma cells using radioimmunoprecipitation assay buffer. Protein concentrations were measured using the BCA protein quantification kit (Thermo Scientific, United States), and 20–50 μg protein extracts were subjected to SDS-PAGE. Then, the proteins were transferred to a nitrocellulose membrane, blocked with 5% nonfat milk, and incubated with primary antibodies for 1 h at room temperature. After incubation with secondary antibodies against mouse (1:10,000) or rabbit (1: 10,000) for 1 h at 4°C overnight, the membrane was visualized using the ECL Western Blotting Detection Kit (Bio-Rad, United States).

### In vitro binding assay of GRP78 and preS1

According to standard protocols, recombinant GST-preS1 was produced in Escherichia coli BL21 cells and purified using glutathione sepharose 4B (GE Healthcare, United States). Ten micrograms of GST or GST fusion proteins and cell lysates were incubated at 4 °C overnight. Supernatants were collected as input, and the sepharose beads were then extensively washed six times with lysis buffer and eluted.

### Immunofluorescence assays

To analyze the colocalization of preS1 and GRP78, we used the preS1 monoclonal antibody and GRP78 monoclonal antibody (Abcam, England). Cells cultured in 48-well plates were washed three times with precooled phosphate-buffered saline, fixed with 4% paraformaldehyde for 10 min, and permeabilized for 10 min at RT with 0.5% Triton X-100. After incubation for 1 h with 3% bovine serum albumin to block nonspecific binding, primary antibodies were added and incubated for 1 h at 37 °C. The bound antibodies were visualized by incubation with secondary antibodies (Alexa Fluor 488 donkey anti-mouse IgG or Alexa Fluor 594 anti-mouse IgG). Images were acquired using a fluorescence microscope.

### Co-immunoprecipitation

Total protein lysates were extracted from hepatoma cells using the IP lysate buffer. The lysate was mixed with 40 μL protein G agarose (Millipore, United States) to avoid nonspecific binding for 2 h at 4 °C. The supernatant was incubated with the primary antibody (Sigma) for 4 h at 4 °C. Subsequently, the mixture was incubated with 60 μL protein agarose for 2 h at 4 °C. The agarose was then washed three times with PBST buffer, boiled, and then loaded onto SDS-PAGE gel for analysis.

### Isothermal titration calorimetry (ITC)

Affinity constants under equilibrium (Ka) were obtained using a Nano ITC instrument. GRP78 protein solution (300 μL) was titrated with repeated injections of 40 μL solution until saturation at 25 °C. Nano Analyze software was used for the integration of heat signals and nonlinear regression analysis of the data.

### Enzyme-linked immunosorbent assay (ELISA)

The expression levels of HBVe antigen (HBeAg) and HBV surface antigen (HBsAg) were measured using an HBeAg/HBsAg ELISA kit (Kehua, China) according to the manufacturer’s instructions. Data were obtained from at least three independent experiments.

### Quantification and differential expression analysis of transcripts

RNA was isolated from HepG2.2.15 cells infected with adenovirus (pAd-GRP78 or pAd-GFP) using TRIzol reagent (Invitrogen, The United States) according to the manufacturer’s instructions, followed by purification on a RNeasy column (Qiagen, German) with DNase treatment. The RNA sequencing library was processed using the Ion Total RNA-Seq Kit v2 (Life Technologies, United States). Then, each library preparation was sequenced using the Ion PI™ chip and raw reads were created. The clean sequences were mapped to the reference genome using Misplacing software, and the gene expression level was normalized to the reads per kilobase per million. To identify differentially expressed genes (DEGs), the DEseq package was used to filter the DEGs for the HepG2.2.15-pAdGRP78 and HepG2.2.15-pAd-GFP groups. After statistical analysis, we screened the DEGs with a fold change > 1.5 or fold change < 0.667, and the false discovery rate threshold was < 0.05. Gene Ontology (GO) analysis was performed using the GO Term Finder tool to identify the main function of the DEGs. To identify significantly enriched pathways, pathway annotation was performed using the Kyoto Encyclopedia of Genes and Genomes database. The significantly enriched pathways were computed based on Fisher’s test of the hypergeometric distribution at p < 0.05.

### Phage display

Phage panning using GRP78 as the target was conducted as previously described with minor modifications (47). The prokaryotic expression and purified GRP78 were diluted to 100 μg/mL in the Tris buffered saline (50 mM Tris, 150 mM NaCl, pH 7.5), and 150 μL of protein was adsorbed onto a polystyrene microtiter plate. It was then coated with GRP78, incubated for 16 h at 4 °C, and blocked (2 h at 37 °C) with Tris buffered saline containing 5 mg/mL bovine serum albumin. Then, a phage library (New England Biolabs, USA) was added. After three rounds of phage panning, a single clone was isolated for DNA sequencing.

### Molecular docking

To calculate the binding energies between GRP78 and the peptide molecule, we performed molecular docking to screen the orientation pattern of the peptides in the binding groove of the GRP78 molecules and residues involved in the interaction. The molecular structure of GRP78 was downloaded from the RCSB Protein Data Bank (code: 6HAB. http://www.rcsb.org/structure/6HAB) and the docking analysis was performed by AutoDock4.2 (http://autodock.scripps.edu/). The binding model with the best energy score was selected for subsequent analysis. The images for clarifying the interactions between GRP78 and the peptide were generated using Pymol (https://pymol.org/2/).

### Microscale thermophoresis (MST)

The interaction between GRP78 and preS1 was confirmed by microscale thermophoresis (48). Experiments were performed with an MST power of 60%, LED power of 40%, and capillaries with a hydrophobic coating under standard conditions. Purified GRP78-His and preS1-GST (preS1-p1-GST, preS1-p2-GST, and preS1-p3-GST) was buffer exchanged into phosphate-buffered saline buffer (pH 7.4), and its concentration was adjusted to 10 μM using UV absorbance. GRP78-His was fluorescently labeled with NT-647 following the manufacturer’s protocol. The preS1-GST solution was diluted to a concentration range of 1.0 × 10-10 to 1.0 × 10-3 mM, and the preS1-GST fusion protein was incubated with the labeled protein for 10 min in the interaction buffer. The samples were then loaded into Monolith NT.115 Capillaries (NanoTemper Technologies) using 50% IR laser power and an LED excitation source (λ= 470 nm) at room temperature. The Kd values were calculated for the interactions between GRP78-His and preS1-GST using NanoTemper Analysis 1.2.20 software. The measurements were replicated three times.

### Statistical analysis

Statistical analysis. All the data are represented as means±standard errors (SE). Significance was determined using a Student’s t test, and standard error was calculated from independent experiments.

## ACKNOWLEDGMENTS

This work was supported by National Science and Technology Major Project (2017ZX10202203, 2018ZX10732202), the National Natural Science Foundation of China (81661148057), Municipal Natural Science Foundation of Chongqing (cstc2018jscx-msyb0004, cstc2020jscx-fyzxX0020).

We thank Haitao Guo (Cancer Virology Program, UPMC Hillman Cancer Center, University of Pittsburgh School of Medicine, Pittsburgh, PA, United States of America) for valuable suggestions and Prof. Ninshao Xia for providing the HepG2-NTPC cells and HBV transgenic mice.

Conceptualization, D.W. and A. H.; Methodology, Y. S., X. J., S. W., J. L. H. Z. and X.F.; Investigation, Y. S., X. J., S. W., J. L. Y. Y., X. Z. and J. W.; Formal Analysis, H. P., M. L., H. Z. and H. Z.; Writing-Original Draft, Y. S.; Writing-Review & Editing, D.W.; Funding Acquisition, D.W. and A. H.; Supervision, D.W. and A. H.

We declare no competing interests.

